# One-carbon metabolism is required for epigenetic stability in the mouse placenta

**DOI:** 10.1101/2023.04.04.535221

**Authors:** Claire E. Senner, Ziqi Dong, Miguel R. Branco, Erica D. Watson

**Affiliations:** Centre for Trophoblast Research, University of Cambridge, Cambridge UK; Department of Physiology, Development, and Neuroscience, University of Cambridge, Cambridge UK; Centre for Genomics and Child Health, Blizard Institute, Faculty of Medicine and Dentistry, Queen Mary University of London, London UK

## Abstract

One-carbon metabolism, including the folate cycle, has a crucial role in fetal development though its molecular function is complex and unclear. The hypomorphic *Mtrr^gt^* allele is known to disrupt one-carbon metabolism, and thus methyl group availability, leading to several developmental phenotypes (e.g., neural tube closure defects, fetal growth anomalies). Remarkably, previous studies showed that some of the phenotypes were transgenerationally inherited. Here, we explored the genome-wide epigenetic impact of one-carbon metabolism in placentas associated with fetal growth phenotypes and determined whether specific DNA methylation changes were inherited. Firstly, methylome analysis of *Mtrr^gt/gt^* homozygous placentas revealed genome-wide epigenetic instability. Several DMRs were identified including at the *Cxcl1* gene promoter and at the *En2* gene locus, which may have phenotypic implications. Importantly, we discovered hypomethylation and ectopic expression of a subset of ERV elements throughout the genome of *Mtrr^gt/gt^* placentas with broad implications for genomic stability. Next, we determined that sperm DMRs in males from the *Mtrr^gt^* model were reprogrammed in the placenta with little evidence of direct or transgenerational germline DMR inheritance. However, some sperm DMRs were associated with placental gene misexpression despite normalisation of DNA methylation, suggesting the inheritance of an alternative epigenetic mechanism. Integration of published histone ChIP-seq datasets with sperm methylome and placenta transcriptome data from the *Mtrr^gt^* model point towards H3K4me3 deposition at key loci suggesting that histone modifications might play a role in epigenetic inheritance in this context. This study sheds light on the mechanistic complexities of one-carbon metabolism in development and epigenetic inheritance.

## Introduction

It is well established that the vitamin folate (also known as folic acid) is important for fetal development. A highly recognisable example is increased risk of neural tube closure defects (e.g., spina bifida) in babies that result from maternal dietary folate deficiency [1]. In fact, folic acid supplementation during pregnancy and folate fortification programmes improves pregnancy outcomes [2, 3]. Beyond the neural tube, other developmental defects (e.g., fetal growth restriction [4, 5], congenital heart defects [6]) and pregnancy disorders [7, 8] are associated with dietary deficiency and/or mutations in key enzymes involved in its metabolism. Although well studied, the molecular role of folate metabolism during development is complex and not well understood. One-carbon metabolism, which includes the folate and methionine cycles, is required by all cells for thymidine synthesis and for methyl groups involved in a broad range of methylation reactions [9]. As a result, it is hypothesised that rapidly proliferating cells in a developing fetus and placenta requires one-carbon metabolism for DNA synthesis and general epigenetic regulation. The specific epigenomic targets of one-carbon metabolism that drive developmental phenotypes remain unclear.

To explore the specific molecular role of one-carbon metabolism during development, we study a mouse model with a hypomorphic mutation in the *Mtrr* (methionine synthase reductase) gene through a gene-trap (gt) insertion [10]. During one-carbon metabolism, folate metabolites are required to transmit methyl groups for the methylation of homocysteine by methionine synthase (MTR) to form methionine and tetrahydrofolate [11]. Methionine acts as precursor for S-adenosylmethionine (SAM), which in turn serves as the sole methyl-donor for substrates involved in epigenetic regulation (e.g., DNA, histones, RNA) among other substrates [12]. Importantly, MTRR activates MTR through the reductive methylation of its vitamin B_12_ co-factor [13–15]. The hypomorphic *Mtrr^gt^* mutation reduces *Mtrr* transcript expression to a level that is sufficient to diminish MTR activity by 60% of controls [10, 13]. Consequently, the progression of one-carbon metabolism is disrupted by the *Mtrr^gt^* mutation as evidenced by plasma hyperhomocysteinemia [10, 13] and widespread changes in DNA methylation patterns [10, 16, 17]. Additionally, *Mtrr^gt/gt^* mice display several phenotypes similar to the clinical features of folate deficiency in humans [18] or human *MTRR* mutations [19, 20] including macrocytic anemia [21] and neural tube closure defects (NTDs) [10, 22]. Beyond this, other phenotypes have emerged in *Mtrr^gt/gt^* mice reflecting a broader influence of impaired one-carbon metabolism on development. These phenotypes include fetal growth defects (such as fetal growth restriction (FGR), fetal growth enhancement (FGE), or developmental delay) [10, 23], complications during implantation (e.g., twinning, skewed implantation) [10, 22], haemorrhages, and/or congenital malformations (such as congenital heart defects and poor placentation) [10, 22, 24]. Therefore, the *Mtrr^gt^* mouse model is ideal for exploring the molecular consequences of defective one-carbon metabolism during growth and development.

Remarkably, the *Mtrr^gt^* mouse line is also a unique mammalian model of transgenerational epigenetic inheritance that occurs via the maternal grandparental lineage [10, 17]. Through highly controlled genetic pedigrees and embryo transfer experiments, we previously showed that an *Mtrr^+/gt^* genotype in male or female mice (i.e., the F0 generation) initiates multigenerational inheritance of developmental phenotypes in their wildtype (*Mtrr^+/+^*) grandprogeny (i.e., the F2-F4 generations) [10, 23]. In general, the mechanism of multigenerational epigenetic inheritance is not well understood. In the context of the *Mtrr^gt^* mouse line, we hypothesise that alterations in the epigenome of the F0 germline caused by abnormal one-carbon metabolism is inherited by the wildtype offspring of the next generation (and potentially beyond) to influence gene expression during development [10, 17, 25]. Given the role of MTRR in one-carbon metabolism, and thus in cellular methylation, we initially focused on DNA methylation patterns. Through a targeted analysis, we previously determined that developmental phenotypes at E10.5 caused by an intrinsic or ancestral *Mtrr^gt^* mutation were associated with locus-specific changes in DNA methylation linked to gene misexpression [10, 17]. The effect was particularly striking in the placenta at key genes involved in the regulation of fetal growth and metabolism, even in wildtype placentas derived from an *Mtrr^+/gt^*ancestor [10]. It is also clear that epigenetic instability of DNA methylation occurs in mature spermatozoa of *Mtrr^gt^* males [17] displaying otherwise normal spermatogenesis and sperm function [26]. However, the extent to which these altered germline methylation patterns are recapitulated in (or inherited by) the somatic cells of the progeny and grandprogeny is currently not well understood in the *Mtrr^gt^*mouse line.

In this study, we investigate the whole placental methylomes of mice exposed to intrinsic (e.g., *Mtrr^gt/gt^*) or ancestral (e.g., F2 *Mtrr^+/+^*) impaired one-carbon metabolism to explore the breadth and specific genomic locations of DNA methylation changes. We relate the specific changes in DNA methylation to gene expression over a spectrum of fetal growth (i.e., FGR or FGE) to investigate whether the epigenetic changes are functionally important. Lastly, by comparing sperm and placenta methylome datasets from *Mtrr^gt^* mice, we aim to better understand whether germline methylation changes that result from abnormal one-carbon metabolism are reprogrammed and/or inherited by the next generation. Ultimately, these analyses delve into the mechanistic complexities of one-carbon metabolism and epigenetic inheritance.

## Results

### Global analysis of the *Mtrr^gt^*^/*gt*^ placental methylome

First, we analysed the extent to which impaired one-carbon metabolism affected the placental methylome and ascertained whether there was an impact on fetal growth. Our initial focus was on *Mtrr^gt^*^/*gt*^ placentas of conceptuses derived from *Mtrr^gt/gt^* intercrosses (Figure 1A). We carried out high-throughput sequencing of immunoprecipitated methylated DNA (meDIP-seq) from C57Bl/6J control and *Mtrr^gt^*^/*gt*^ placentas at E10.5. *Mtrr^gt/gt^* placentas were divided into two phenotypic groups including those from fetuses that were phenotypically normal (PN) or were FGR based on crown-rump length [10, 22]. Placentas from C57Bl/6J mice were controls since the *Mtrr^gt^* allele was backcrossed into the C57Bl/6J genetic background [10]. However, we previously identified four regions of structural variation between C57Bl/6J and the *Mtrr^gt^* line [17]. To avoid false discovery of changes in DNA methylation during the meDIP-seq data analysis, these regions were excluded bioinformatically along with the 20 Mb region of 129P2Ola/Hsd genomic sequence surrounding the gene-trapped *Mtrr* allele [16, 17], that remained after eight backcrosses [10].

**Figure 1.**
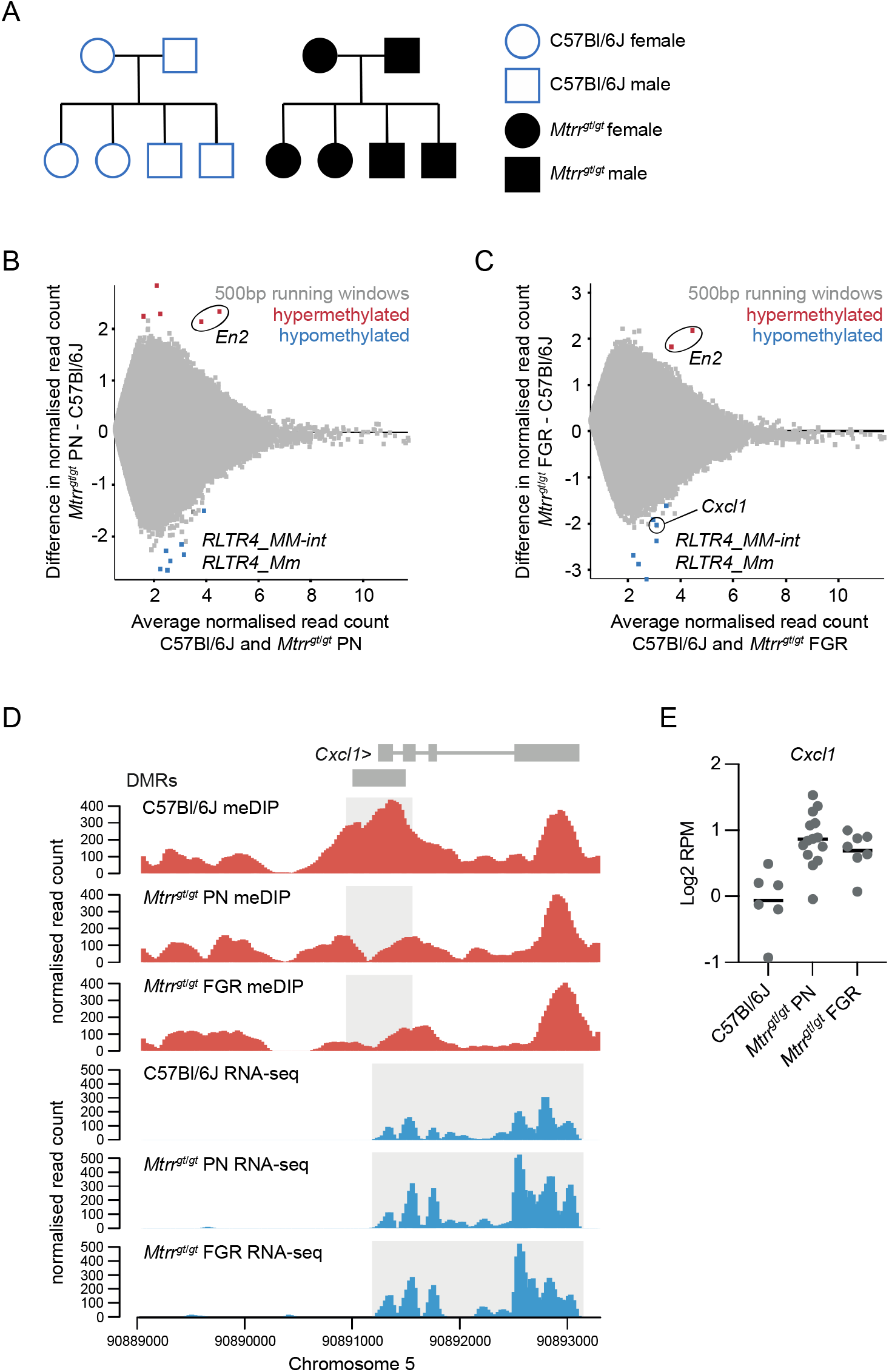
Analysis of the *Mtrr^gt/gt^* placental methylome. (A) Schematised C57Bl6J control pedigree (blue outline, white fill) and *Mtrr^gt^*^/*gt*^ pedigree (black outline, black fill) used in this study. Square, males; Circles, females. **(B-C)** MA plot of *log_2_* normalised meDIP-seq read counts of 500bp contiguous regions in (B) C57Bl/6J placentas and *Mtrr^gt^*^/*gt*^ placentas from phenotypically normal (PN) fetuses, and (C) C57Bl/6J placentas and *Mtrr^gt^*^/*gt*^ placentas associated with fetal growth restriction (FGR). Hypermethylated (red) and hypomethylated (blue) differentially methylated 500 bp regions (DMR) were identified using EdgeR. **(D)** Data tracks showing normalised meDIP-seq (red) and RNA-seq (blue) reads across the *Cxcl1* locus on mouse chromosome 5 in C57Bl/6J, *Mtrr^gt^*^/*gt*^ PN and *Mtrr^gt^*^/*gt*^ FGR placentas. DMR and transcript expression are highlighted in light grey. **(E)** Graph showing *Cxcl1* transcript expression (*log_2_*RPM) ascertained by RNA-seq, in C57Bl/6J, *Mtrr^gt^*^/*gt*^ PN, and *Mtrr^gt^*^/*gt*^ FGR placentas at E10.5. In all cases data was normalised to the largest data store. For meDIP-seq: C57Bl/6J, N=8 placentas; *Mtrr^gt/gt^* PN, N=7 placentas; *Mtrr^gt/gt^* FGR, N=7 placentas. For RNA-seq: C57Bl/6J, N=6 placentas; *Mtrr^gt/gt^* PN, N=14 placentas; *Mtrr^gt/gt^* FGR, N=7 placentas.

At the global level, we found that the distribution of meDIP-seq reads across different genomic features were not significantly different between C57Bl/6J control and *Mtrr^gt^*^/*gt*^ placentas even when phenotypic severity was considered (Figure S1A). Furthermore, meDIP-seq datasets from individual placentas did not cluster by *Mtrr* genotype or fetal growth phenotype when data store similarity tools were implemented (Figure S1B). As global DNA methylation patterns were similar between experimental groups, we next ascertained differences in DNA methylation at individual loci compared to control placentas. DMRs were defined using the EdgeR function embedded within Seqmonk software (www.bioinformatics.babraham.ac.uk) with default settings (p<0.05, with multiple testing correction) assessing 500 bp contiguous regions. The resulting DMRs were further filtered for regions that displayed a *log_2_*fold change (FC) >1 in DNA methylation compared to controls. Only a few DMRs were present in *Mtrr^gt^*^/*gt*^ placentas (i.e., PN: 9 DMRs; FGR: 13 DMRs), though both hyper- and hypomethylated regions were observed (Figure 1B-C, Supplementary data file 1). The low number of DMRs caused by the *Mtrr^gt^* allele suggested that DNA methylation changes were subtle regardless of fetal growth phenotypes or were ‘hidden’ by our analysis of whole placentas as individual cell types might be differently affected.

Despite the low number of total DMRs, three key findings emerged (explored further below). Firstly, only one placental DMR associated with the misexpression of a protein-coding gene (*Cxcl1*; Figure 1D-E). Secondly, we identified two hypermethylated DMRs in *Mtrr^gt^*^/*gt*^ placentas that were common to PN and FGR conceptuses with potential developmental importance (Figure 1B-C). These two 500 bp regions were contiguous and represented one single 1 kb DMR within the single intron of the *En2* gene (Figure 2A). This DMR was also previously identified in mature sperm and embryos at E10.5 from *Mtrr^gt^*^/*gt*^ mice [17] and was flagged since it might play a role in inheritance. Lastly, of the 12 hypomethylated DMRs that were identified, ten were associated with endogenous retroviruses (ERVs). Strikingly, the majority of these DMRs (7/10) overlapped with ERV1 elements of the RLTR4 subclass (Figure 1B-C) indicating potential implications for genetic stability. Further exploration into the potential importance of these three findings was explored below.

**Figure 2.**
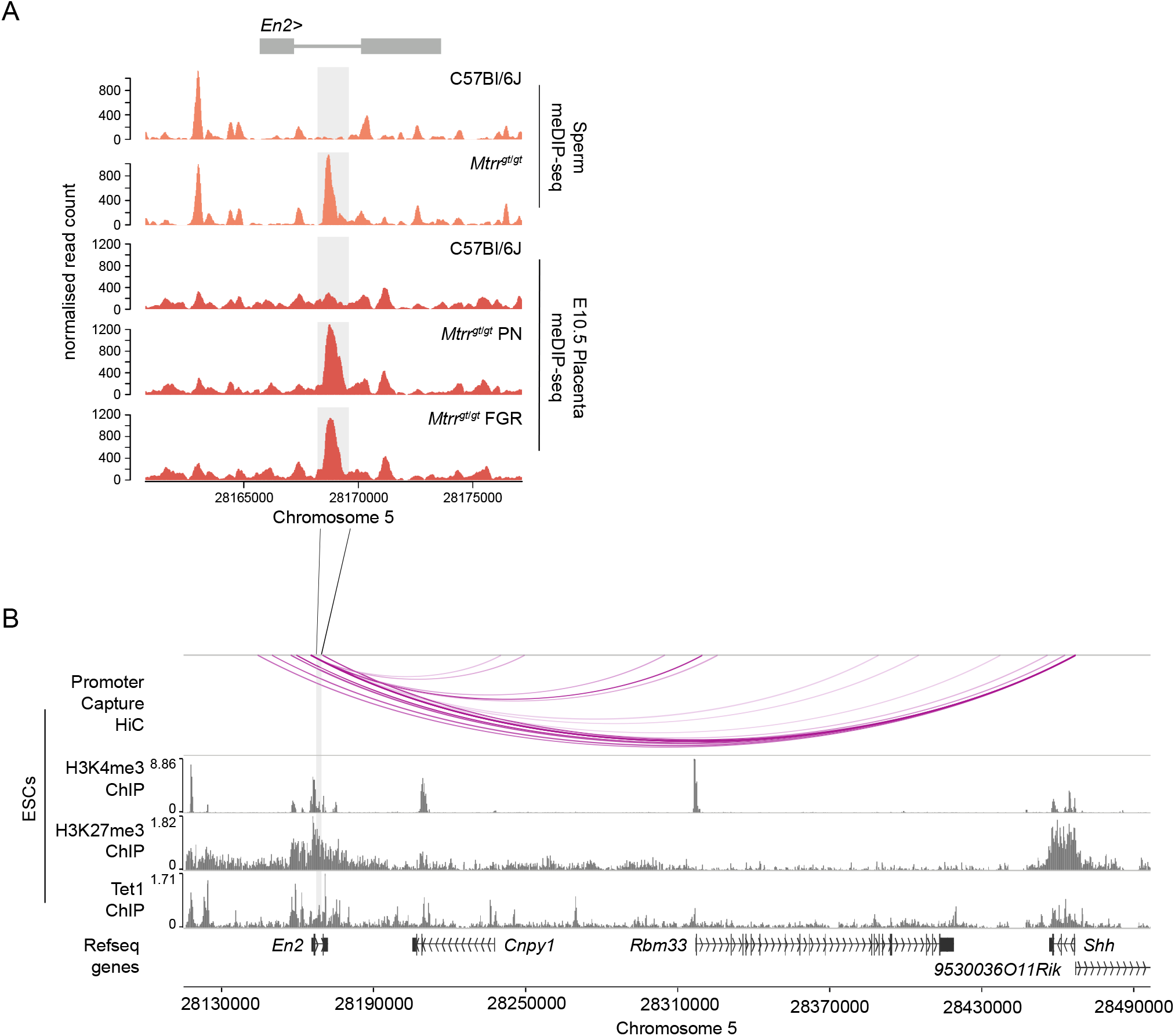
*En2* DMR as a potential regulator of developmentally important genes in embryonic lineage. **(A)** Data tracks showing normalised meDIP-seq read counts across the *En2* gene in sperm from C57BL/6J and *Mtrr^gt^*^/*gt*^ mice (orange) and placentas from C57Bl/6J and *Mtrr^gt^*^/*gt*^ conceptuses at E10.5 (red). Placentas were associated with either phenotypically normal (PN) or fetal growth restricted (FGR) fetuses. The *En2* DMR is highlighted in light grey. **(B)** Data tracks showing promoter capture Hi-C-based interactions (purple lines) alongside H3K27me3, H3K4me3 and Tet1 ChIP-seq peaks in wildtype mouse embryonic stem cells (ESCs) at the *En2* locus and in downstream genes. See Supplementary Data File 3 for data sources. For sperm meDIP-seq: C57Bl/6J, N=8 males; *Mtrr^gt/gt^*, N=8 males. For placenta meDIP-seq: C57Bl/6J, N=8 placentas; *Mtrr^gt/gt^* PN, N=7 placentas; *Mtrr^gt/gt^*FGR, N=7 placentas.

### Potential canonical regulation of placental *Cxcl1* expression by DNA methylation

To explore whether altered placental DNA methylation caused by the *Mtrr^gt/gt^* genotype had a gene regulatory effect, we employed published RNA-seq datasets from *Mtrr^gt/gt^* placentas at E10.5 associated with PN and FGR fetuses [27]. Using DESeq, the RNA-seq data was assessed for differentially expressed genes that were within 2 kb of a DMR (identified in *Mtrr^gt/gt^* placentas) and had transcript levels with a *log_2_*FC>0.6 compared to control placentas. The *Cxcl1* gene (chemokine (C-X-C motif) ligand 1) was the only dysregulated gene identified in this context. CXCL1 is important for decidual angiogenesis to promote maternal blood flow into the implantation site [28].

We observed that hypomethylation at a DMR located in the promoter of *Cxcl1* was associated with a modest upregulation of *Cxcl1* transcripts (Figure 1D, 1E). This finding exemplifies canonical regulation of a gene by DNA methylation. Since *Mtrr^gt/gt^* placentas from both PN and FGR fetuses displayed hypomethylation of the *Cxcl1* DMR and upregulation of *Cxcl1* transcripts, these molecular changes were likely insufficient to drive the fetal growth phenotype.

### *En2* DMR as a potential regulator of developmentally important genes

Given the prevalence of the *En2* DMR (Figure 2A) in mature sperm, embryos and placentas of *Mtrr^gt/gt^* mice [17], its potential functional importance was explored. The *En2* gene encodes a homeobox transcription factor that, when knocked out in mice, leads to autism-spectrum disease-like behaviours [29–31] that are accompanied by cerebellar foliation defects [32] and loss of GABAergic interneurons in somatosensory and visual cortical areas [33, 34]. Indeed, *En2* mRNA is expressed in multiple regions of the developing brain [35] and is involved in neurogenesis [36].

Low levels of *En2* transcripts were reported by RNA-seq in the ectoplacental cone [37] (a population of trophoblast progenitor cells in the mouse placenta). However, our RNA-seq data from whole C57Bl/6J control placentas at E10.5 [27] showed that *En2* transcripts were very lowly expressed (Figure S2) and thus, *En2* might be considered as an unexpressed gene in the placenta at this developmental stage. Importantly, hypermethylation of the *En2* DMR in *Mtrr^gt/gt^* placentas was not associated with a change in *En2* transcript levels (Figure S2) indicating that the *En2* DMR is an unlikely regulator of *En2* gene expression in this context.

To investigate a broader regulatory role of the *En2* DMR, the histone methylation landscape was explored for potential interactions between the DMR and promoter regions of neighbouring genes. To do this, we assessed published H3K4me3 and H3K27me3 ChIP-seq datasets and promoter capture Hi-C datasets from wildtype mouse trophoblast stem cells (TSCs) [38]. TSCs are an *in vitro* model of undifferentiated trophoblast cells of the placenta [39], and these datasets represent the most suitable available for analysis. No enrichment of H3K4me3 or H3K27me3 modifications was evident at the *En2* DMR in TSCs nor were any DMR-promoter interactions apparent in the immediate genomic region (Figure S2). Accordingly, genes downstream of the *En2* DMR were also expressed at normal levels in *Mtrr^gt/gt^* placentas indicating that hypermethylation of this region had little to no effect on cis regulation of gene expression in the placenta (Figure S2).

We previously showed that the *En2* DMR was also hypermethylated in whole *Mtrr^gt/gt^*embryos at E10.5 as determined by bisulfite pyrosequencing [17]. Therefore, we explored additional ChIP-seq and promoter capture Hi-C datasets in wildtype mouse embryonic stem cells (ESCs) [38] to determine whether the *En2* DMR had regulatory potential in the embryonic lineage. We observed that the *En2* DMR had hallmarks of a regulatory locus in ESCs since it was bivalently marked by the enrichment of repressive H3K27me3 and active H3K4me3 modifications (Figure S2) in a manner that poises this region for activation upon cell differentiation [40]. The genomic region defined by the *En2* DMR in ESCs was also enriched for the DNA demethylating enzyme TET1 (Figure S2), which typically co-localises with polycomb complexes and contributes to keeping unmethylated enhancers and promoters methylation-free [41]. Furthermore, promoter capture Hi-C experiments in ESCs [38] revealed a potential interaction of the *En2* DMR with the promoters of nearby genes, including *Cnpy1* (canopy FGF signalling regulator 1), *Rbm33* (RNA binding motif 33), and the developmental regulator *Shh* (sonic hedgehog) (Figure S2). Given these data, we hypothesised that specific ectopic hypermethylation of the *En2* DMR in *Mtrr^gt/gt^* embryos might affect expression of surrounding genes with developmental consequences. Further analysis of the developmental role of the *En2* DMR in the embryo is required, particularly in the context of abnormal one-carbon metabolism.

### Hypomethylation and ectopic expression of ERVs indicates epigenetic instability in the ***Mtrr^gt^* mouse line**

We identified several hypomethylated DMRs in *Mtrr^gt^*^/*gt*^ placentas at E10.5 that were associated with ERV1 elements of the RLTR4 subfamily (separately annotated as *RLTR4_Mm* and *RLTR4_MM-int* for the LTRs and internal region, respectively; Figure 1B, 1C). RLTR4 elements are relatively young retrotransposons that are closely related to murine leukemia virus and that, at least in some mouse strains, remain transpositionally active [42]. Some transposable elements (e.g., IAPs) are highly methylated and resistant to epigenetic reprogramming to avoid genomic transposition [43]. It is unclear whether this is the case for RLTR4 elements. The RLTR4 elements associated with placental DMRs in this study typically displayed a pro-viral, full-length configuration, rather than being solo LTRs or other isolated fragments. Furthermore, six out of seven of the DMRs mapped to two discrete genomic loci on mouse chromosomes 11 and 18, including in regions that were intragenic (and antisense) to *Camk2b* and upstream of the gene *Pik3c3*, respectively (Figure 3A, 3B). Remarkably, a loss of DNA methylation at these DMRs corresponded with ectopic expression of the RLTR4 element in *Mtrr^gt/gt^* placentas, independent of the fetal growth phenotype (Figure 3A, 3B). No expression changes in the associated protein-coding genes were observed (Figure 3A, B). Therefore, the genomic regions demarcated by these DMRs appear to require methylation to repress the ERV element activity and not to regulate cis gene expression in the placenta.

**Figure 3.**
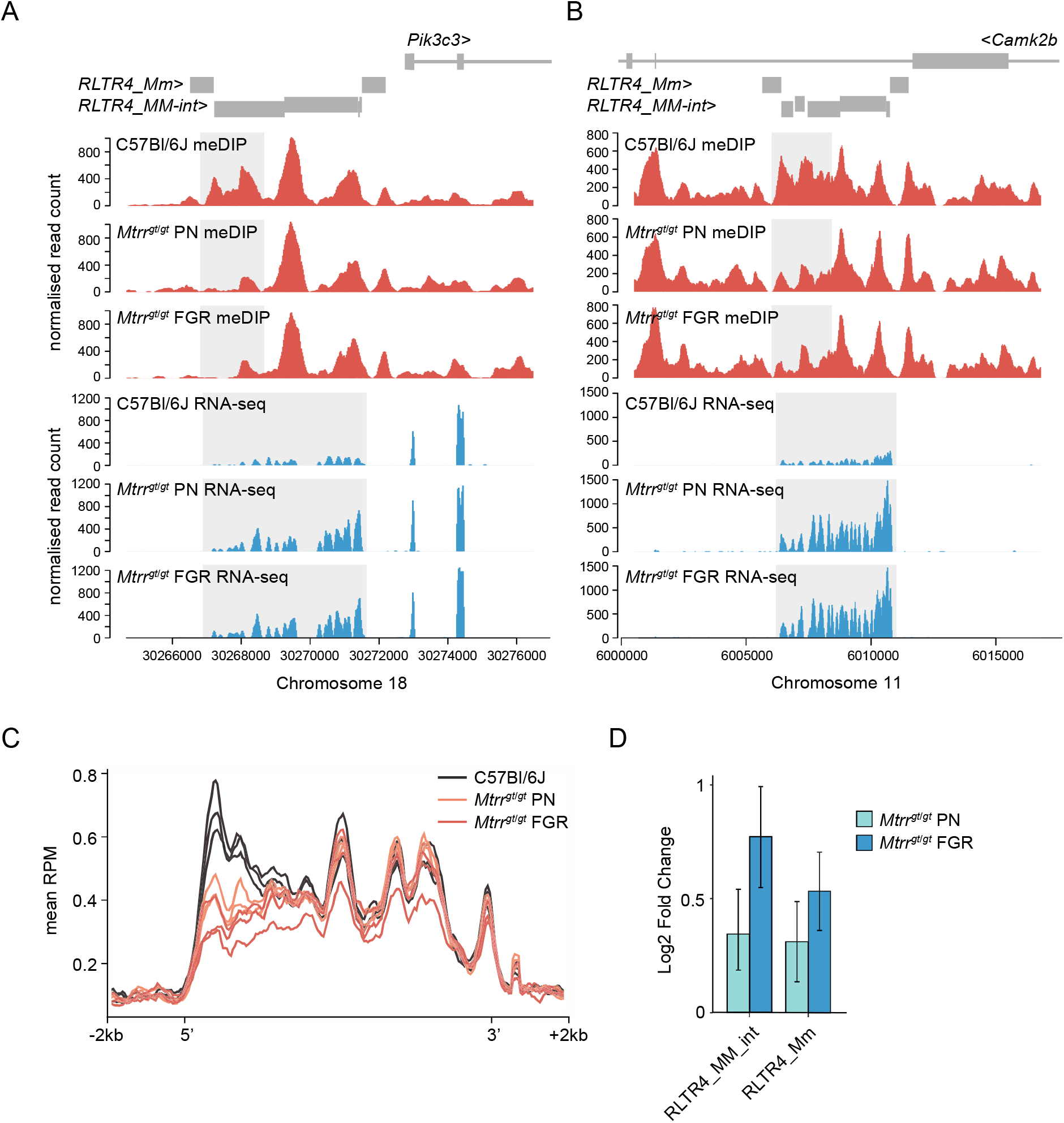
Analysis of DNA methylation and transcript expression at ERVs in *Mtrr^gt/gt^* placentas at E10.5. (**A and B**) Data tracks showing normalised meDIP-seq (red) and RNA-seq (blue) reads across full-length ERVs comprising *RLTR4_Mm* and *RLTR4_MM-int* elements on mouse **(A)** chromosome 18 associated with the *Pik3c3* gene and **(B)** chromosome 11 associated with the *Camk2b* gene in placentas of C57Bl/6J and *Mtrr^gt^*^/*gt*^ conceptuses at E10.5. Placentas from phenotypically normal (PN) and fetal growth restricted (FGR) fetuses were assessed. DMR and transcript expression are highlighted in light grey. **(C)** Graph representing the average meDIP-seq reads mapping across all full-length RLTR4 elements +/-2kb in the genome in individual placentas from C57Bl/6J (black), *Mtrr^gt/gt^* PN (orange) and *Mtrr^gt/gt^* FGR (red) fetuses at E10.5. **(D)** Enrichment of *RLTR4_Mm* and *RLTR4_MM-int* expression in placentas from *Mtrr^gt^*^/*gt*^ PN (light blue) and *Mtrr^gt^*^/*gt*^ FGR (dark blue) fetuses at E10.5 relative to C57Bl/6J control placentas as determined by RNA-seq. For meDIP-seq: C57Bl/6J, N=8 placentas; *Mtrr^gt/gt^* PN, N=7 placentas; *Mtrr^gt/gt^* FGR, N=7 placentas. For RNA-seq: C57Bl/6J, N=6 placentas; *Mtrr^gt/gt^* PN, N=14 placentas; *Mtrr^gt/gt^* FGR, N=7 placentas.

Due to their repetitive nature and evolutionary young age, the mapability of short sequencing reads to RLTR4 elements is low. Therefore, to fully appreciated the dysregulation of DNA methylation at RLTR4 elements in *Mtrr^gt/gt^* placentas, the placental meDIP-seq (this study) and RNA-seq [27] datasets from C57Bl/6J and *Mtrr^gt/gt^*placentas at E10.5 were re-mapped to include non-unique reads by using random assignment (bowtie2) for meDIP-seq data and an expectation-maximisation algorithm (SQuIRE [44]) for RNA-seq data. When considered globally, the remapped data revealed consistent DNA hypomethylation at the 5’ end of RLTR4 full-length elements in *Mtrr^gt^*^/*gt*^ placentas at E10.5 (Figure 3C). This pattern of DNA hypomethylation was associated with transcript enrichment of global *RLTR4_Mm* and *RLTR4_MM-int* elements in *Mtrr^gt^*^/*gt*^ placentas compared to controls (Figure 3D). Differential expression analysis of individual elements uncovered significant upregulation of twenty-four *RLTR4_Mm* or *RLTR4_MM-int* elements that converged upon 15 full-length loci (Supplementary Data File 2). None of these methylation changes associated with altered expression of nearby protein-coding genes in *Mtrr^gt/gt^* placentas. Furthermore, methylation and transcriptional dysregulation at *RLTR4_Mm* and *RLTR4_Mm-int* elements was unlikely to regulate fetal growth since the RLTR4 elements were similarly affected in *Mtrr^gt/gt^* placentas associated with PN and FGR fetuses (Figure 3D). Overall, these data reinforced the hypothesis that epigenetic instability is inherent to the *Mtrr^gt^* mouse line [10, 17] with implications for genetic stability and phenotype establishment beyond FGR.

### *Mtrr^gt^*^/^*^gt^* sperm DMRs are reprogrammed in the placenta but correspond with placental gene misexpression

In our previous study of epigenetic inheritance in the *Mtrr^gt^* mouse line [17], we found that a small number of candidate DMRs identified in sperm were not recapitulated in embryos or placentas at E10.5 when interrogated by bisulfite pyrosequencing. Here, we validated this finding on a genome-wide scale by comparing our sperm [17] and placenta meDIP-seq datasets from control and *Mtrr^gt/gt^* mice. First, we harmonised DMR calling between datasets by reanalysing the sperm meDIP-seq datasets according to our analysis of the placenta meDIP-seq data. Hypermethylated sperm DMRs in *Mtrr^gt/gt^* males that were previously located in very highly methylated regions in control sperm and described as false positives [17], were also clearly identifiable by the current analysis. Accordingly, we screened out these DMRs using whole genome bisulphite sequencing data to quantify absolute methylation levels across all DMRs [45]. Similar to our candidate-based approach [17], DNA methylation patterns in nearly all genomic regions identified as sperm DMRs in *Mtrr^gt/gt^* males were normal in *Mtrr^gt/gt^* placentas at E10.5 compared to control placentas (Figure 4A- C). This finding occurred regardless of the *Mtrr^gt/gt^* fetal growth phenotype. As expected, the only exception was the common hypermethylated *En2* DMR that appeared in both sperm and placentas (Figure 4B-C). Conversely, the placental DMRs that overlap with RLTR4 elements were normally methylated in sperm of *Mtrr^gt/gt^* males (relative to control sperm) suggesting that germline transposon silencing is maintained and unlikely to play a key role in epigenetic inheritance mechanisms in the *Mtrr^gt^* model. Overall, these results indicated that placenta and sperm DMRs in *Mtrr^gt^*^/*gt*^ mice were tissue-specific and that most of the sperm DMRs were effectively reprogrammed in the pre-implantation embryo or during placental development, notwithstanding the shared *Mtrr^gt/gt^* genotype of the parental and offspring generations. These data might negate DNA methylation as a mechanistic factor in epigenetic inheritance within the *Mtrr^gt^* model. Instead, an alternative (or additional) mechanism of abnormal one-carbon metabolism might influence heritability of other epigenetic factors, such as histone modifications or small non-coding RNA content in germ cells.

**Figure 4.**
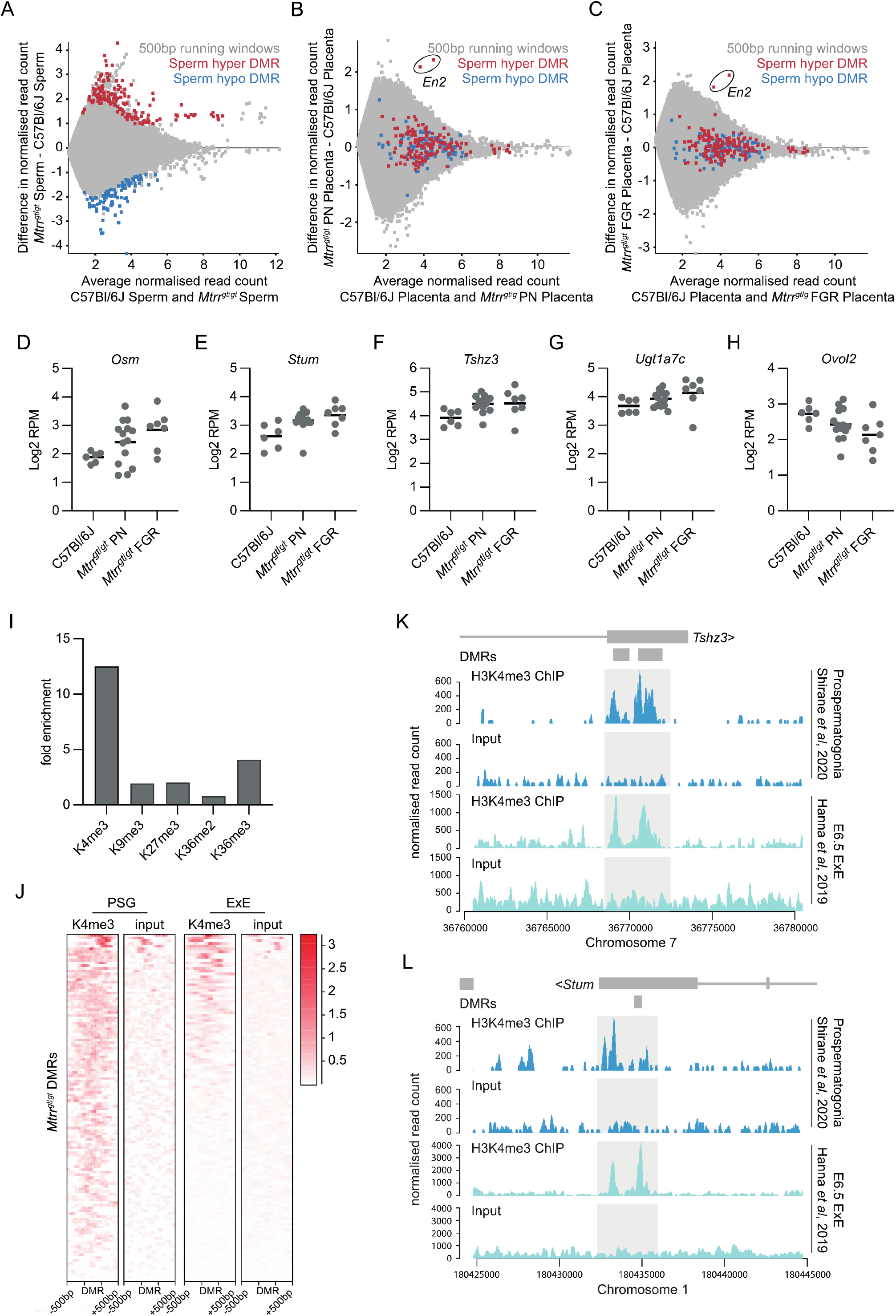
Sperm DMRs in *Mtrr^gt/gt^* males were normalised in *Mtrr^gt/gt^* placentas yet associated with transcriptional dysregulation. (A) MA plot of *log_2_* normalised meDIP-seq read counts of 500 bp contiguous regions in sperm from C57Bl/6J and *Mtrr^gt^*^/*gt*^ males. Hypermethylated (red) and hypomethylated (blue) differentially methylated 500 bp regions (DMR) were identified using EdgeR. (B) MA plot of *log_2_* normalised meDIP-seq read counts of 500 bp contiguous regions in placentas from C57Bl/6J and *Mtrr^gt^*^/*gt*^ phenotypically normal fetuses at E10.5. The genomic regions where sperm DMRs from *Mtrr^gt/gt^* males were identified are highlighted on the placenta data. Hypermethylated sperm DMRs (red), hypomethylated sperm DMRs (blue). The *En2* DMRs are indicated. (C) MA plot of *log_2_* normalised meDIP-seq read counts of 500 bp contiguous regions in placentas from C57Bl/6J fetuses and *Mtrr^gt^*^/*gt*^ fetal growth restricted (FGR) fetuses at E10.5. The genomic regions where sperm DMRs from *Mtrr^gt/gt^* males were identified are highlighted on the placenta data. Hypermethylated sperm DMRs (red), hypomethylated sperm DMRs (blue). The *En2* DMRs are indicated. **(D-H)** Graphs showing placental transcript expression (*log_2_*RPM) of genes that were associated with sperm DMRs including **(D)** *Osm*, **(E)** *Stum*, **(F)** *Tshz3*, **(G)** *Ugt1a7c*, and **(H)** *Ovol2*. Data was ascertained by RNA-seq of placentas from C57Bl/6J and *Mtrr^gt^*^/*gt*^ conceptuses at E10.5. Placentas from phenotypically normal (PN) and fetal growth restricted (FGR) fetuses were assessed. **(I)** Enrichment for specific histone modifications in wildtype prospermatogonia ascertained by ChIP-seq at the 500 bp regions defined as DMRs in sperm of *Mtrr^gt^*^/*gt*^ males. Enrichment determined relative to the baseline genome. **(J)** Probe alignment plot showing H3K4me3 enrichment ascertained by ChIP-seq from wildtype prospermatogonia and extraembryonic ectoderm (ExE) at E6.5 compared to input controls in regions identified as sperm DMRs (+/- 500 bp) in *Mtrr^gt^*^/*gt*^ males. **(K)** and **(L)** Data tracks showing normalised H3K4me3 ChIP-seq reads and input controls for prospermatogonia (dark blue) and extraembryonic ectoderm (ExE) at E6.5 (light blue) in the regions surround the **(K)** *Tshz3* and **(L)** *Stum* sperm DMRs from *Mtrr^gt/gt^* males. Light grey boxes highlight H3K4me3 peaks. Small dark grey boxes indicate the DMRs. See Supplementary Data File 3 for data sources. For sperm meDIP-seq: C57Bl/6J, N=8 males; *Mtrr^gt/gt^*, N=8 males. For placenta meDIP-seq: C57Bl/6J, N=8 placentas; *Mtrr^gt/gt^* PN, N=7 placentas; *Mtrr^gt/gt^* FGR, N=7 placentas. For RNA-seq: C57Bl/6J, N=6 placentas; *Mtrr^gt/gt^* PN, N=14 placentas; *Mtrr^gt/gt^* FGR, N=7 placentas.

Previously, our locus-specific analysis showed that some genes associated with sperm DMRs were misexpressed in somatic tissues despite reprogramming of DNA methylation at these sites [17]. Therefore, we questioned the extent to which this association occurred in the wider placental genome. Using DESeq, the *Mtrr^gt/gt^* placental RNA-seq dataset [27] was assessed for differentially expressed genes that were within 2 kb of a sperm DMR from *Mtrr^gt/gt^* males and had transcript levels with a *log_2_*FC>0.6 compared to control placentas. Five misexpressed genes (i.e., *Stum* (mechanosensory transducer mediator; membrane protein)*, Tshz3* (teashirt zinc finger family member 3; transcription factor)*, Ovol2* (ovo like zinc finger 2; transcription factor), *Osm* (oncostatin m; cytokine), and *Ugt1a7c* (UDP glucouronosyltransferase 1 family, polypeptide A7C; enzyme in glucouronidation pathway) met these criteria but only in *Mtrr^gt/gt^* placentas with FGR (Figure 4D-H). The occurrence of transcriptional disruption despite normal DNA methylation reinforced our hypothesis that abnormal one-carbon metabolism influences other epigenetic mechanisms.

Next, we explored whether histone modifications were present in the developing germline at regions demarcated by sperm DMRs to better understand the broader epigenetic context. To do this, ChIP-seq datasets were analysed for histone mark enrichment in developing wildtype sperm (i.e., prospermatogonia) [46] at specific genomic regions defined by sperm DMRs from *Mtrr^gt/gt^* males. First, we found that 103 out of 252 DMRs (40.9%) overlapped with an H3K4me3 peak in wildtype prospermatogonia. This represented a 12-fold enrichment compared to the baseline genome (i.e., only 3.3% of 500 bp regions across the whole genome overlapped with H3K4me3 peaks) and was substantially more enriched than the other histone modifications at the same locations (Figure 4I). Since H3K4me3 is typically located at gene promoters and indicates transcriptional activation [47], it was an ideal candidate to further explore as an underlying inherited epigenetic mark associated with transcriptional disruption in the placenta. Therefore, we assessed whether wildtype trophoblast progenitor cells at E6.5 (i.e., extraembryonic ectoderm) displayed H3K4me3 enrichment at genomic locations identified as sperm DMRs using a published ChIP-seq dataset [48]. Indeed, a substantial subset of these genomic regions was also enriched for H3K4me3 in extraembryonic ectoderm (Figure 4J). Remarkably, four out of five dysregulated genes in *Mtrr^gt/gt^* placentas that were associated with a sperm DMR in *Mtrr^gt/gt^* males (i.e., *Stum*, *Tshz3*, *Ovol2*, *Ugta7c*) were among those enriched for H3K4me3 in both prospermatogonia and extraembryonic ectoderm (Figure 4K-4L and S3). We might infer from these data that the *Mtrr^gt^* mutation potentially disrupts sperm histone marks, such as H3K4me3, leading to altered patterns of the same histone mark in the early conceptus with implications for gene regulation. Future mechanistic experiments should focus on multigenerational patterns of H3K4me3 in the *Mtrr^gt^* mouse line.

### Sperm DMRs caused by an *Mtrr^gt^* allele were not multi-generationally inherited

Lastly, a transgenerational link was explored in the *Mtrr^gt^* mouse line between sperm DMRs and the placental methylome and transcriptome. To do this, the following matings were performed (Figure 5A): F0 *Mtrr^+/gt^* males were mated with C57Bl/6J control females, and the resulting F1 *Mtrr^+/+^* females were selected for mating with C57Bl/6J males to generate F2 *Mtrr^+/+^* conceptuses. F2 *Mtrr^+/+^* conceptuses were rigorously phenotyped at E10.5 [10], and the placentas from PN and FGE fetuses were examined. Previous directed locus-specific analyses in F2 *Mtrr^+/+^* placentas by bisulfite pyrosequencing indicated significant alteration of DNA methylation patterns caused by an ancestral *Mtrr^gt^* allele [10]. Here, the broader methylome of whole F2 *Mtrr^+/+^* placentas was assessed via meDIP-seq. When compared with C57Bl/6J controls, there were no significant differences in the distribution of meDIP reads across genomic features (Figure S1C) and no clustering of biological replicates according to phenotype or pedigree (Figure S1D) indicating similar global methylation among experimental groups. Few DMRs were identified by meDIP-seq analysis including one hypermethylated DMR in F2 *Mtrr^+/+^* placentas from PN fetuses (Figure 5B) and 11 DMRs in FGE-associated F2 *Mtrr^+/+^* placentas (Figure 5C). Closer analysis revealed that nine of the hypomethylated DMRs from F2 *Mtrr^+/+^* FGE placentas were clustered in two locations on chromosomes 14 and 17, which have been frequently susceptible to mapping artefacts in our datasets and so were excluded. The remaining three placental DMRs from F2 *Mtrr^+/+^* placentas were in nondescript genomic regions (Supplementary Data file 1). Importantly, the RLTR4 elements identified in *Mtrr^gt/gt^* placentas exhibited normal levels of DNA methylation and transcript expression in the F2 *Mtrr*^+/+^ placentas relative to control placentas (Figure S4). This result suggested that the changes in DNA methylation described in *Mtrr^gt^*^/*gt*^ placentas are intrinsically associated with the *Mtrr^gt^* allele and are unlikely to be transgenerationally inherited or caused by genetic differences between the C57Bl/6J control and *Mtrr^gt^* mouse lines.

**Figure 5.**
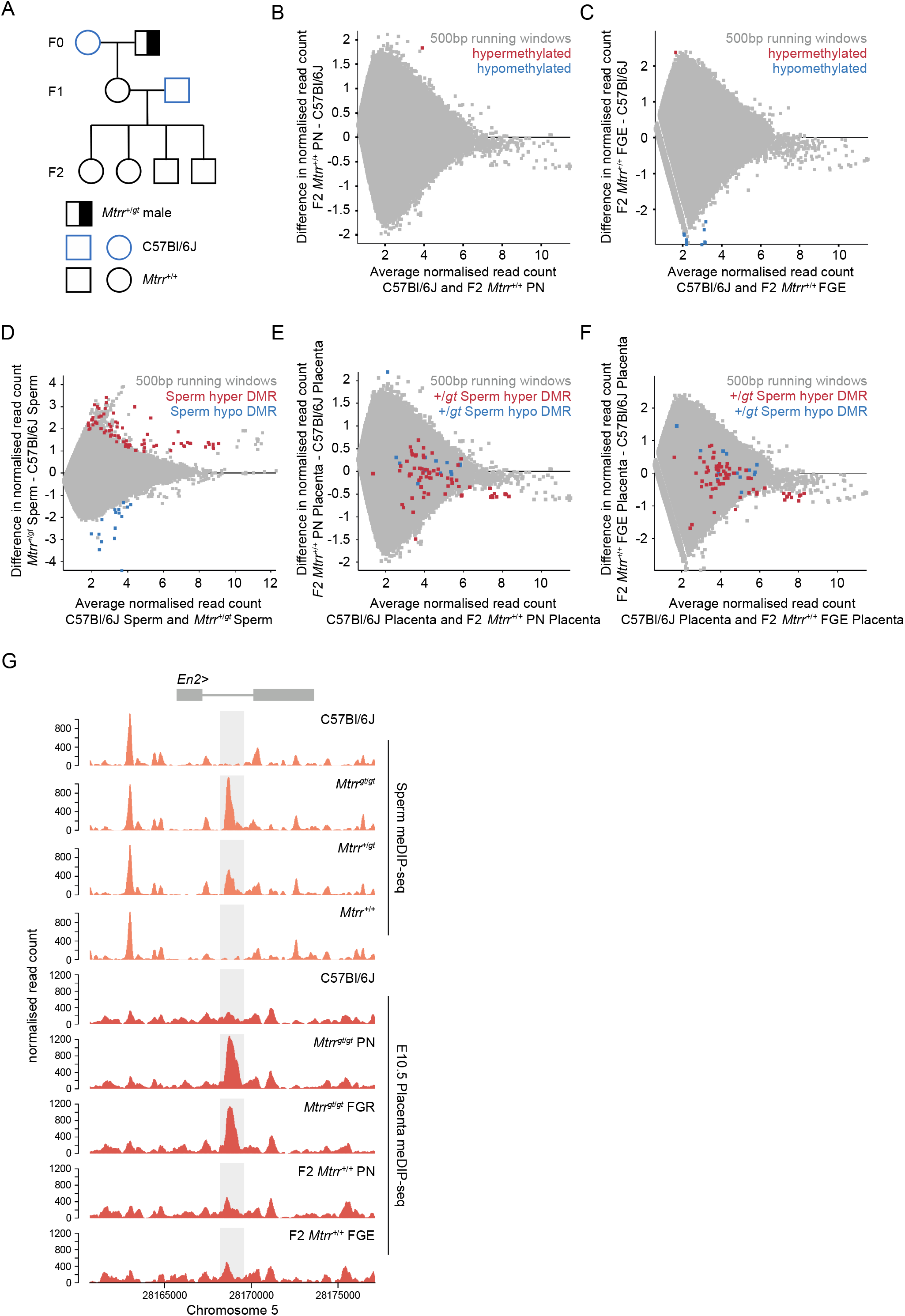
Sperm DNA methylation patterns were not transgenerationally inherited in the *Mtrr^gt^*mouse line. (A) *Mtrr*^+/gt^ maternal grandfather pedigree used in this study. Squares, males; circles females; blue outline, C57Bl/6J mouse line; black outline, *Mtrr^gt^* mouse line; white fill, *Mtrr^+/+^*; half black/half white fill; *Mtrr^+/gt^*. F0, parental generation; F1, first filial generation; F2, second filial generation. (B) MA plot of *log_2_* normalised meDIP-seq read counts of 500 bp contiguous regions in placentas at E10.5 from C57Bl/6J conceptuses and F2 *Mtrr*^+/+^ conceptuses derived from F0 *Mtrr^+/gt^* maternal grandfathers. Placentas from phenotypically normal (PN) fetuses were assessed. Hypermethylated (red) and hypomethylated (blue) differentially methylated regions (DMRs) were determined relative to control placentas using EdgeR. (C) MA plot of *log_2_* normalised meDIP-seq read counts of 500 bp contiguous regions in placentas at E10.5 from C57Bl/6J conceptuses and F2 *Mtrr*^+/+^ fetal growth enhanced (FGE) conceptuses derived from F0 *Mtrr^+/gt^* maternal grandfathers. Hypermethylated (red) and hypomethylated (blue) DMRs were determined using EdgeR. (D) MA plot of *log_2_* normalised meDIP-seq read counts of 500 bp contiguous regions in sperm from C57Bl/6J and F0 *Mtr^r+/gt^* males. Hypermethylated DMRs (red), hypomethylated DMRs (blue). (E) MA plot of *log_2_* normalised meDIP-seq read counts of 500 bp contiguous regions in placentas from C57Bl/6J and F2 *Mtrr^+/+^* PN fetuses at E10.5. The genomic regions where sperm DMRs from F0 *Mtrr^+/gt^* males were identified are highlighted on the placenta data. Hypermethylated sperm DMRs (red), hypomethylated sperm DMRs (blue). (F) MA plot of *log_2_* normalised meDIP-seq read counts of 500 bp contiguous regions in placentas at E10.5 from C57Bl/6J fetuses and F2 *Mtrr^+/+^* FGE fetuses. The genomic regions where sperm DMRs from F0 *Mtrr^+/gt^* males were identified are highlighted on the placenta data. Hypermethylated sperm DMRs (red), hypomethylated sperm DMRs (blue). (G) Data tracks across the *En2* gene showing normalised meDIP-seq read counts in sperm (orange) from C57BL/6J males and *Mtrr^gt^*^/*gt*^, *Mtrr*^+/*gt*^ and *Mtrr*^+/+^ males together with meDIP read counts in placentas at E10.5 (red) associated with C57Bl/6J fetuses, *Mtrr^gt^*^/*gt*^ PN and FGR fetuses, and F2 *Mtrr*^+/+^ PN and FGE fetuses. The *En2* DMR is highlighted in light grey. In all cases data was normalised to the largest data store. For sperm meDIP-seq: C57Bl/6J, N=8 males, F0 *Mtrr^+/gt^*, N=8 males. For placenta meDIP-seq: C57Bl/6J, N=8 placentas; F2 *Mtrr*^+/+^ PN, N=8 placentas; F2 *Mtrr*^+/+^ FGE, N=3 placentas.

When the meDIP-seq datasets from sperm of F0 *Mtrr^+/gt^* males (Figure 5D) [17] were compared to placentas of F2 *Mtrr^+/+^*conceptuses (PN and FGE) at E10.5, there was no DMR overlap (Figure 5E, 5F). This finding reinforces our hypothesis that DMRs in the *Mtrr^gt^* mouse line are not inherited from germline to somatic cells over multiple generations. This was even the case at the *En2* DMR, which was present in sperm from F0 *Mtrr^+^*^/*gt*^ males and not in F2 *Mtrr^+/+^* placentas (Figure 5G). In this context, our sperm and placenta methylome data revealed that the dosage of the *Mtrr^gt^* allele in mice correlated with the degree of hypermethylation at the *En2* DMR (Figure 5G). Therefore, the *En2* locus was particularly responsive to *Mtrr*-driven disruption of one-carbon metabolism. Altogether, these data further separate the transmission of differential methylation via the germline from fetal growth phenotype inheritance in the *Mtrr^gt^* mouse line.

## Discussion

Despite its well-studied role in development and disease, the molecular function of one-carbon metabolism is complex and not well understood. Here, we used the *Mtrr^gt^* mouse model to explore the epigenetic role of one-carbon metabolism by assessing the placental methylome in association with fetal growth phenotypes. In doing so, we identified several genomic regions in *Mtrr^gt/gt^* placentas with altered DNA methylation including in a gene promoter that conceivably regulates *Cxcl1* gene expression in a canonical manner, in a presumptive developmental regulatory region located within the *En2* gene, and in a subset of RLTR4 transposable elements. While unlikely to underlie the fetal growth phenotypes, it is possible that these DNA methylation changes are functionally relevant in other tissue types and/or for driving other phenotypes. For instance, CXC chemokine expression from peripheral blood mononuclear cells correlates with folate and homocysteine levels in human subjects [49]. Alternatively, knocking out the mouse gene *Mtfhr* to disrupt folate metabolism causes cerebellar patterning defects that are associated with down- regulation of *En2* gene expression [50]. Ultimately, our findings support widespread epigenetic instability in the *Mtrr^gt^*mouse line.

Our previous locus-specific analyses indicated that placentas exposed to intrinsic or ancestral *Mtrr^gt^*allele are epigenetically unstable [10, 16, 17]. Yet, we identified fewer placenta DMRs by meDIP-seq than were expected despite using standard analysis parameters that yielded many sperm DMRs (e.g., 13 placenta DMRs vs 252 sperm DMRs from *Mtrr^gt/gt^* mice). It is possible that placental DNA methylation is less sensitive than sperm to impaired one-carbon metabolism. DNA in trophoblast cells is globally hypomethylated compared to other cell types [51] and changes in DNA methylation might be less striking in this context. The use of whole placentas that contain multiple cell types (e.g., trophoblast cell subtypes, fetal vascular endothelium, and maternal decidua and immune cells) with their own DNA methylation and transcriptional signatures [52, 53] might confound our analysis to some extent. The placental DMRs that we identified are likely present throughout the tissue, while other undetected DMRs may be confined to a single cell type and not appreciated in our analysis. Assaying the placenta at earlier developmental time points when the trophoblast progenitor population is more homogeneous may be informative.

Alternatively, single cell-based sequencing methods may uncover additional DMRs in *Mtrr^gt/gt^* placentas at E10.5 that correlate with cell-type specific transcriptional dysregulation and phenotypes.

Transposable elements, which make up ∼40% of the mammalian genome [54], are heavily methylated to suppress transposition causing deleterious mutation. In this study, we observed hypomethylation and ectopic expression of several RLTR4 elements in *Mtrr^gt/gt^* placentas, which might have profound consequences to genomic stability during development. While whole genome sequencing revealed that *de novo* mutation rates are similar in control and *Mtrr^gt/gt^* mice [17], it is still possible that increased transposition might occur in this context, generating structural variation with implications for phenotype inheritance. Since sperm from *Mtrr^gt/gt^* males showed normal RLTR4 DNA methylation, we propose that DNA methylation was poorly maintained in early embryogenesis or in placenta development to cause hypomethylation at these sites. Although RLTR4 elements were the only transposons identified by meDIP-seq in this study, DNA methylation patterns of variably methylated intracisternal A particle (VM-IAP) retrotransposons are also considerably shifted in *Mtrr^gt/gt^* mice as determined by bisulfite pyrosequencing [16]. KRAB- ZPFs are known to regulate VM-IAPs [55], and mechanistically, the *Mtrr^gt^* locus contains a cluster of 129P2Ola/Hsd-derived KRAB zinc finger proteins (ZFPs) in an otherwise C57Bl/6J background because of the mutagenesis process [16]. However, the KRAB-ZFP clusters within the *Mtrr* locus do not appear to regulate RLTR4 expression [56]. Others have shown that paternal *Mthfr* deficiency in mice causes hypomethylation of L1Md subfamily of LINE-1 retrotransposons [57]. Altogether, these data highlight the importance of one-carbon metabolism in maintaining epigenetic stability at early developmental stages when deleterious transposition events could have profound consequences.

While the mechanistic understanding of epigenetic inheritance remains in its infancy, several candidate epigenetic factors have been identified (e.g., chromatin modifications, small non- coding RNA content in germ cells) [48]. Our data provides genome-wide evidence that nearly all sperm DMRs caused by the *Mtrr^gt^* mutation were epigenetically reprogrammed in the placenta and were not transgenerationally inherited. This contrasts with another study that demonstrates transgenerational inheritance of directed epimutations of DNA methylation in mouse obesity genes along with an obesity phenotype, despite evidence that these epimutations are reprogrammed in primordial germ cells [58]. The lack of DMR inheritance in the *Mtrr^gt^* mouse line suggests that there are paradigm-specific effects. However, there are clues that sperm DMRs caused by the *Mtrr^gt^*mutation might still play a role in epigenetic inheritance since they are associated locus-specific disruption of transcription in *Mtrr^gt/gt^* placentas (this study) and in F2 *Mtrr^+/+^* embryos and adult livers [17] despite being reprogrammed to normal tissue-specific methylation levels. This association evokes a role for other epigenetic mechanisms aside from DNA methylation in epigenetic inheritance mechanisms. We observed enrichment for the activating H3K4me3 histone mark in developing wildtype sperm and trophoblast specifically at genomic locations of sperm DMRs in *Mtrr^gt/gt^* mice including loci associated with placental gene misexpression. Others have shown a similar association in a mouse model of paternal *Mthfr* deficiency [57]. The most striking data that reinforces a potential role for histone H3K4me in epigenetic inheritance comes from a study whereby mice were fed a folate-deficient diet. The resulting sperm displayed alterations in histone H3K4me3 patterns specifically at developmental genes and putative enhancers, a subset of which were retained in the pre-implantation embryos and were associated with gene misexpression [59]. Therefore, we propose that histone methylation should be explored as an inherited epigenetic mechanism in the *Mtrr^gt^*mouse model.

Overall, this study together with our previously published work [10, 16, 17] indicate that one-carbon metabolism is required for the maintenance of epigenetic stability in the placenta and the germline. The widespread effect on the epigenome provides some explanation towards the complex molecular role of folate metabolism during development. Instability of the epigenome can alter transcriptional pathways and genomic stability, with substantial downstream effects on developmental outcome. Single-cell sequencing technology and a broader analysis of epigenetic mechanisms (e.g., histone marks and chromatin structure) in addition to DNA methylation will enable the identification of complex epigenome-phenotype relationships that persist over multiple generations in context of the *Mtrr^gt^* mouse line.

## Methods

### Ethics statement

This research was regulated under the Animals (Scientific Procedures) Act 1986 Amendment Regulations 2012 following ethical review by the University of Cambridge Animal Welfare and Ethical Review Body.

### Mouse model

*Mtrr^Gt(XG334)Byg^* (MGI:3526159) mouse line, referred to as the *Mtrr^gt^* model, was generated when a β*- geo* gene-trap (gt) vector was inserted into intron 9 of the *Mtrr* gene in 129Ola/Hsd embryonic stem cells (ESCs) [10, 13]. *Mtrr^gt^* ECSs were injected into C57Bl/6J blastocysts and upon germline transmission, the *Mtrr^gt^* allele was backcrossed into the C57Bl/6J genetic background for at least eight generations [10]. *Mtrr^+/+^*and *Mtrr^+/gt^* mice were generated from *Mtrr^+/gt^* intercrosses. *Mtrr^gt/gt^* mice were generated by *Mtrr^gt/gt^* intercrosses. Since the *Mtrr^gt^* allele has a multigenerational effect [10, 16, 17], C57Bl/6J mice from The Jackson Laboratories (www.jaxmice.jax.org) were used as controls and were bred in-house and maintained separately from the *Mtrr^gt^* mouse line. The effects of a maternal grandpaternal *Mtrr^gt^* allele were determined with the following pedigree: F0 *Mtrr^+/gt^*males were mated to C57Bl/6J females. The resulting F1 *Mtrr^+/+^*females were mated to C57Bl/6J males to generate F2 *Mtrr^+/+^*conceptuses. Genotyping for *Mtrr^+^* and *Mtrr^gt^* alleles was performed using PCR on DNA extracted from ear tissue or yolk sac using a three-primer reaction resulting in a wildtype band at 252 bp and a mutant band at 383 bp [10]. Primer sequences: primer *a* (5’- GAGATTGGGTCCCTCTTCCAC), primer *b* (5’-GCTGCGCTTCTGAATCCACAG), and primer *c* (5’-CG ACTTCCGGAGCGGATCTC) [10]. All mice were housed in a temperature-and humidity-controlled environment with a 12 h light-dark cycle. All mice were fed a normal chow diet (Rodent No. 3 chow, Special Diet Services) *ad libitum* from weaning, which included (per kg of diet): 1.6 g choline, 2.73 mg folic acid, 26.8 μg vitamin B_12_, 3.4 g methionine, 51.3 mg zinc.

### Dissections and tissue collection

Noon of the day that the vaginal plug was detected was defined as E0.5. Mice were euthanized by cervical dislocation. Fetuses and placentas were dissected in cold 1x phosphate buffered saline (PBS) at E10.5 using a Zeiss SteReo Discovery V8 microscope, scored for phenotypes (see below), and photographed. Fetuses and placentas were weighed and measured separately and snap frozen in liquid nitrogen (stored at -80°C). Both male and female placentas were assessed since no phenotypic sexual dimorphism was identified at E10.5 [23].

### Phenotyping at E10.5

Conceptuses were rigorously scored for gross phenotypes during dissection and allocated to the phenotypic categories that were previously defined, including phenotypically normal (PN), fetal growth enhancement (FGE), fetal growth restriction (FGR), developmental delay, severe abnormalities (e.g., congenital heart defects, neural tube closure defects, hemorrhages, skewed conceptus orientation, twinning, etc), and resorption [10, 22]. Notably, conceptuses with >1 phenotype were counted once and classified by the most severe phenotype observed. Only PN, FGR, and FGE conceptuses were assessed in this study. Phenotype parameters are defined below.

*PN conceptuses*: fetuses and placentas met all developmental milestones appropriate for the developmental stage according to e-Mouse Atlas Project (https://www.emouseatlas.org/emap/home.html). PN fetuses at E10.5 contained 30-39 somite pairs and had crown-rump lengths (for E10.5) that were within two standard deviations (sd) from the mean of C57Bl/6J fetuses, putting them within the normal range for growth. All PN conceptuses lacked severe abnormalities identified via gross assessment.

*FGR and FGE conceptuses:* Conceptuses with FGR and FGE lacked severe abnormalities identified via gross dissection and met the staging criteria for E10.5 (i.e., 30-39 somite pairs). Yet, the fetuses displayed crown-rump lengths that were ≥ 2 sd below (for FGR) or above (for FGE) the mean measurement for C57Bl/6J fetuses. Conceptus size was unaffected by litter size in all pedigrees and stages assessed [27].

### Methylated DNA immunoprecipitation (meDIP) and next generation sequencing

MeDIP-Seq was carried out as described previously [60]. Briefly, genomic DNA was purified from whole placentas at E10.5 and was sonicated to yield 150–600 bp fragments, and adaptors for paired-end sequencing (Illumina) were ligated using NEBNext DNA Sample Prep Reagent Set 1 (New England Biolabs). Immunoprecipitations were carried out using 500ng DNA per sample, 1.25μg anti-5mC antibody (Eurogentec BI-MECY-0100) and 10μL Dynabeads coupled with M-280 sheep anti-mouse antibody (Invitrogen). Pulled down DNA was amplified for 12 cycles with adapter-specific indexed primers. Final clean-up and size selection was carried out with AMPure-XP SPRI beads (Beckman Coulter). Libraries were quantified and assessed using the Kapa Library Quantification Kit (Kapa Biosystems) and Bioanalyzer 2100 System (Agilent). Indexed libraries were sequenced (50-bp paired-end) on an Illumina HiSeq 2500 sequencer. Raw FASTq data were trimmed with TrimGalore, using default parameters, and unique reads mapped to the *Mus musculus* GRCm38 genome assembly using Bowtie2. Data analysis was carried out using SeqMonk software (www.bioinformatics.babraham.ac.uk).

### Transposable element analyses

To include transposon-derived reads that do not map uniquely, the meDIP-seq datasets were re- aligned using the default settings of bowtie2 to assign reads with multiple equally best alignments to one of those locations at random. Average methylation levels over pro-viral, full-length elements were generated after merging Repeatmasker annotations for RLTR4_Mm and RLTR4_MM-int elements. RNA-seq data was analysed using SQuIRE [44], which assigns multimapping reads using an expectation-maximisation algorithm, and provides both subfamily-level and single copy-level information. Differential expression analysis was performed using SQuIRE’s Call function.

## Data availability

All data are available from the corresponding author upon reasonable request. The source and accession numbers of processed WGBS, ChIP-seq and Hi-C data sets used in this study can be found in Supplementary Data File 3.

## Author Contributions

C.E.S. and E.D.W. conceptualised the study and designed the experiments. E.D.W. performed the dissections and phenotyped the conceptuses. C.E.S. and Z.D. generated the placenta meDIP libraries. C.E.S. and M.B. designed and performed the bioinformatics analyses. C.E.S., M.B., and E.D.W. collected and analysed the data. C.E.S., M.B., and E.D.W. interpreted the results. E.D.W. and C.E.S. wrote the manuscript. All authors read and revised the manuscript.

## Acknowledgements

Next generation sequencing was performed at the Babraham Institute (UK).

## Supplementary figure legends

**Figure S1.**
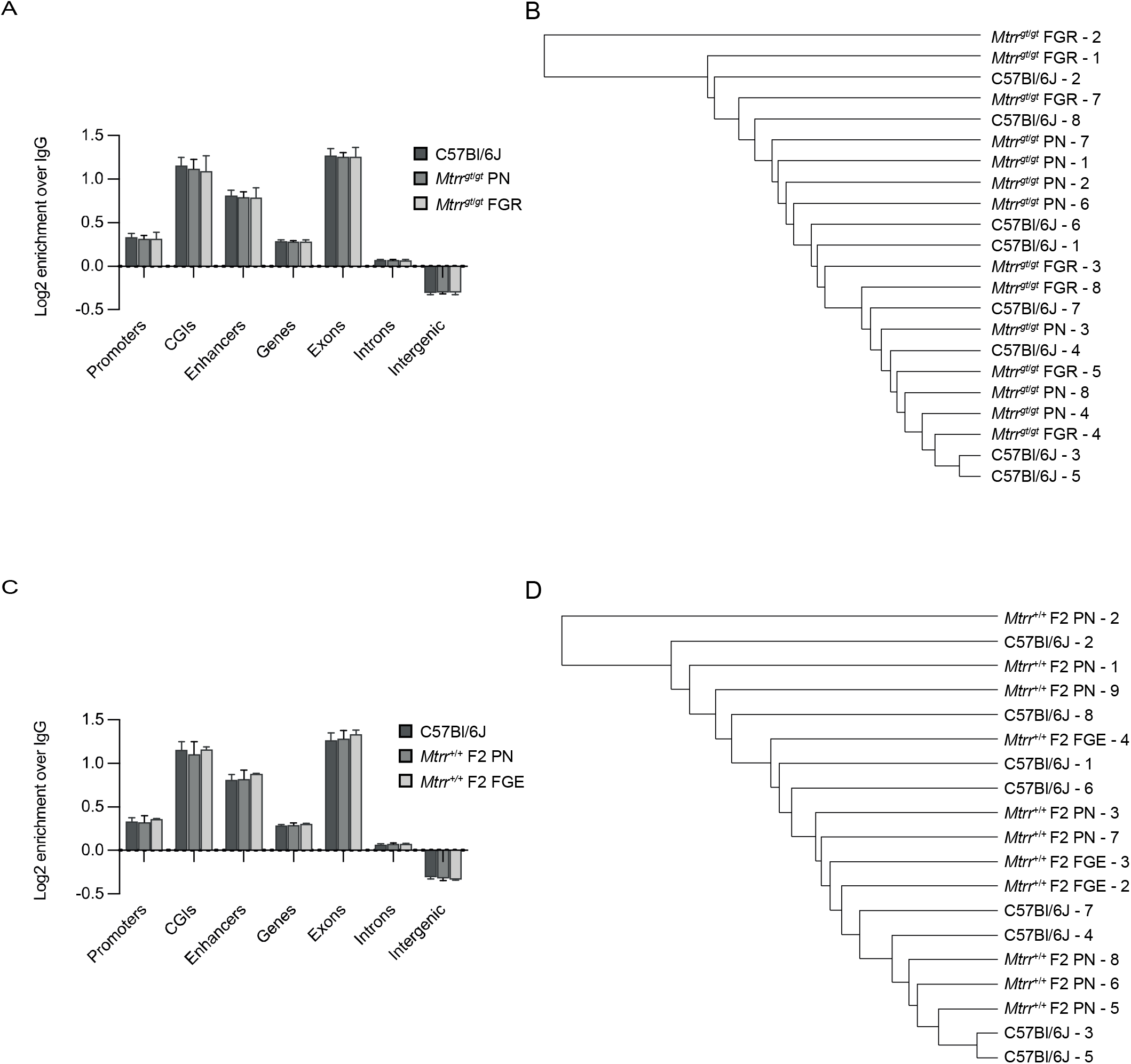
Distribution of meDIP-seq reads across genomic features and phenotypes. (A) Enrichment of meDIP-seq reads (*log_2_* scale) across genomic features relative to IgG in C57Bl/6J placentas, and *Mtrr^gt^*^/*gt*^ placentas from conceptuses that were phenotypically normal (PN) or displayed fetal growth restriction (FGR) at E10.5. (B) Data Store tree displaying all replicates from C57Bl/6J, *Mtrr^gt^*^/*gt*^ PN, and *Mtrr^gt^*^/*gt*^ FGR placentas. Clustering is based on normalised read counts of 500bp contiguous regions tiled over the genome. (C) Enrichment of meDIP-seq reads (*log_2_* scale) across genomic features relative to IgG in C57Bl/6J placentas at E10.5 and F2 *Mtrr*^+/+^ placentas from conceptuses that were PN or displayed fetal growth enhancement (FGE) at E10.5. (D) Data Store tree displaying all replicates from C57Bl/6J, F2 *Mtrr*^+/+^ PN, and F2 *Mtrr*^+/+^ FGE placentas. Clustering is based on normalised read counts of 500bp contiguous regions tiled over the genome. CGIs, CpG islands.

**Figure S2.**
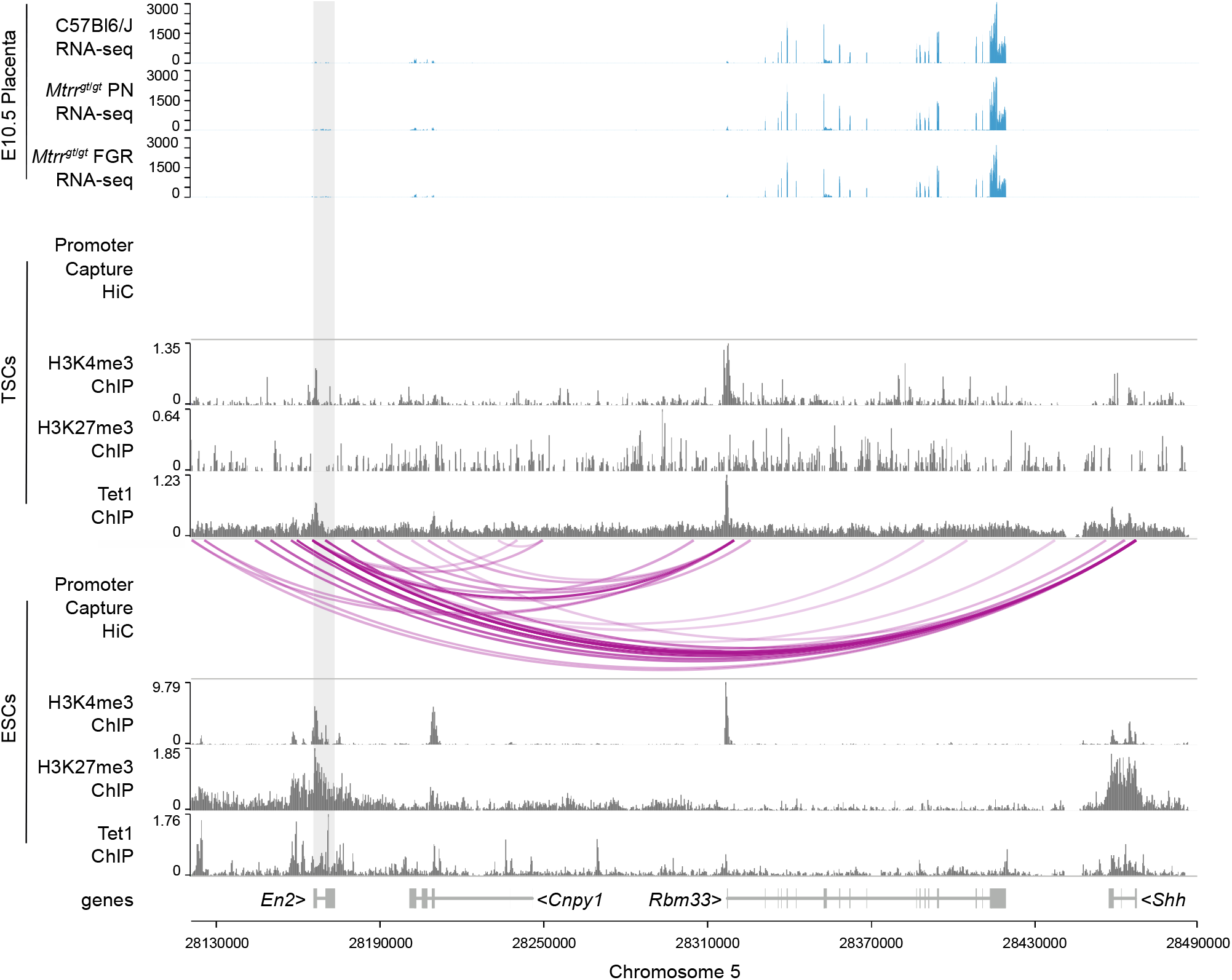
Gene expression and chromatin marks within the genomic region surrounding the mouse *En2* gene in placentas at E10.5, and ESCs and TSCs. (top) RNA-seq data tracks showing gene expression (blue) in the genomic region of the *En2* DMR in placentas at E10.5 from C57Bl/6J and *Mtrr^gt^*^/*gt*^ conceptuses. Placentas from phenotypically normal (PN) and fetal growth restricted (FGR) fetuses were considered. **(middle)** Promoter capture Hi-C based interactions and H3K27me3, H3K4me3 and Tet1 ChIP-seq peaks in wildtype mouse trophoblast stem cells (TSCs). **(bottom)** Promoter capture Hi-C based interactions and H3K27me3, H3K4me3 and Tet1 ChIP-seq peaks in wildtype mouse embryonic stem cells (ESCs). See Supplementary Data File 3 for data sources. For RNA-seq: C57Bl/6J, N=6 placentas; *Mtrr^gt/gt^* PN, N=14 placentas; *Mtrr^gt/gt^* FGR, N=7 placentas.

**Figure S3.**
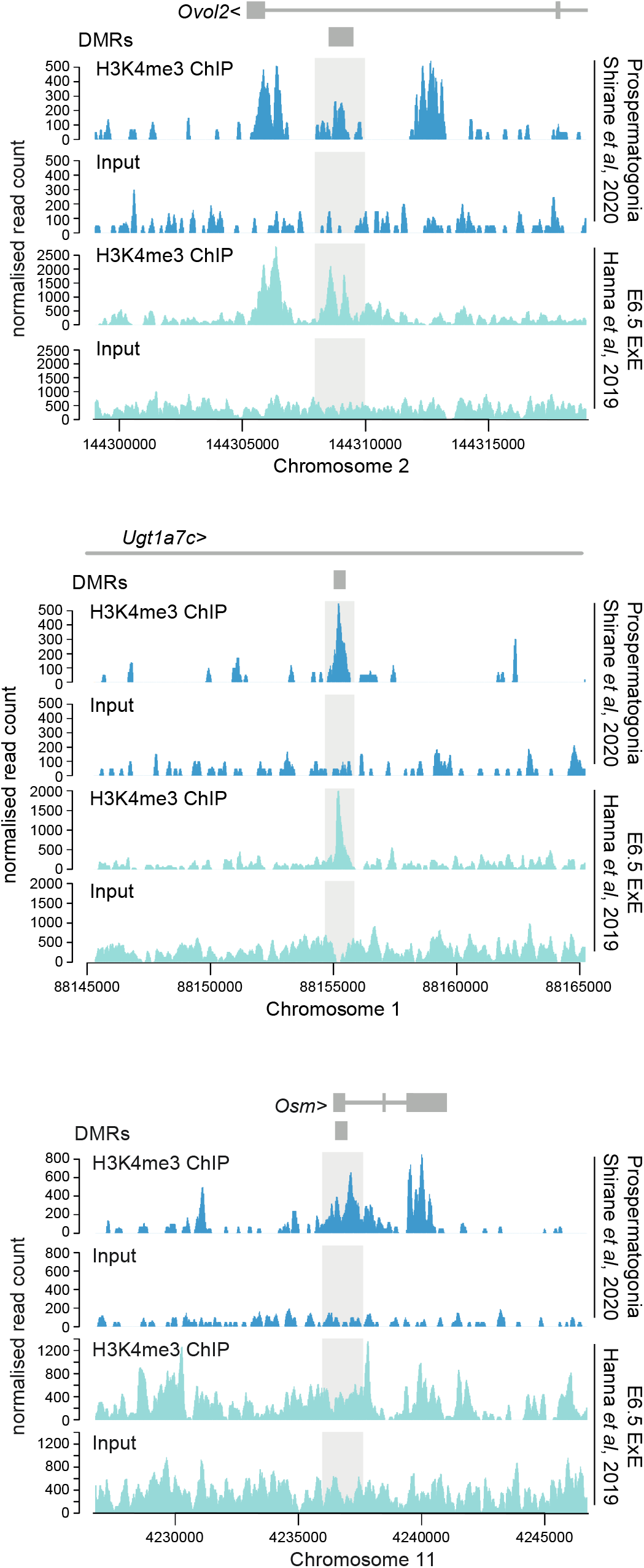
H3K4me3 enrichment in prospermatogonia and extraembryonic ectoderm in regions defined by sperm DMRs and placental gene misexpression in *Mtrr^gt/gt^*mice. Data tracks showing H3K4me3 ChIP-seq reads and input controls for prospermatogonia (dark blue peaks) and extraembryonic ectoderm at E6.5 (light blue peaks). The regions shown are associated with sperm DMRs (dark grey) identified in *Mtrr^gt/gt^* males that correspond to gene misexpression in *Mtrr^gt/gt^* placentas at E10.5 including the *Ovol2* DMR (top), *Ugt1a7* DMR (middle), and *Osm* DMR (bottom). Light grey boxes highlight H3K4me3 peaks within the region specified as a sperm DMR. See Supplementary Data File 3 for data sources.

**Figure S4.**
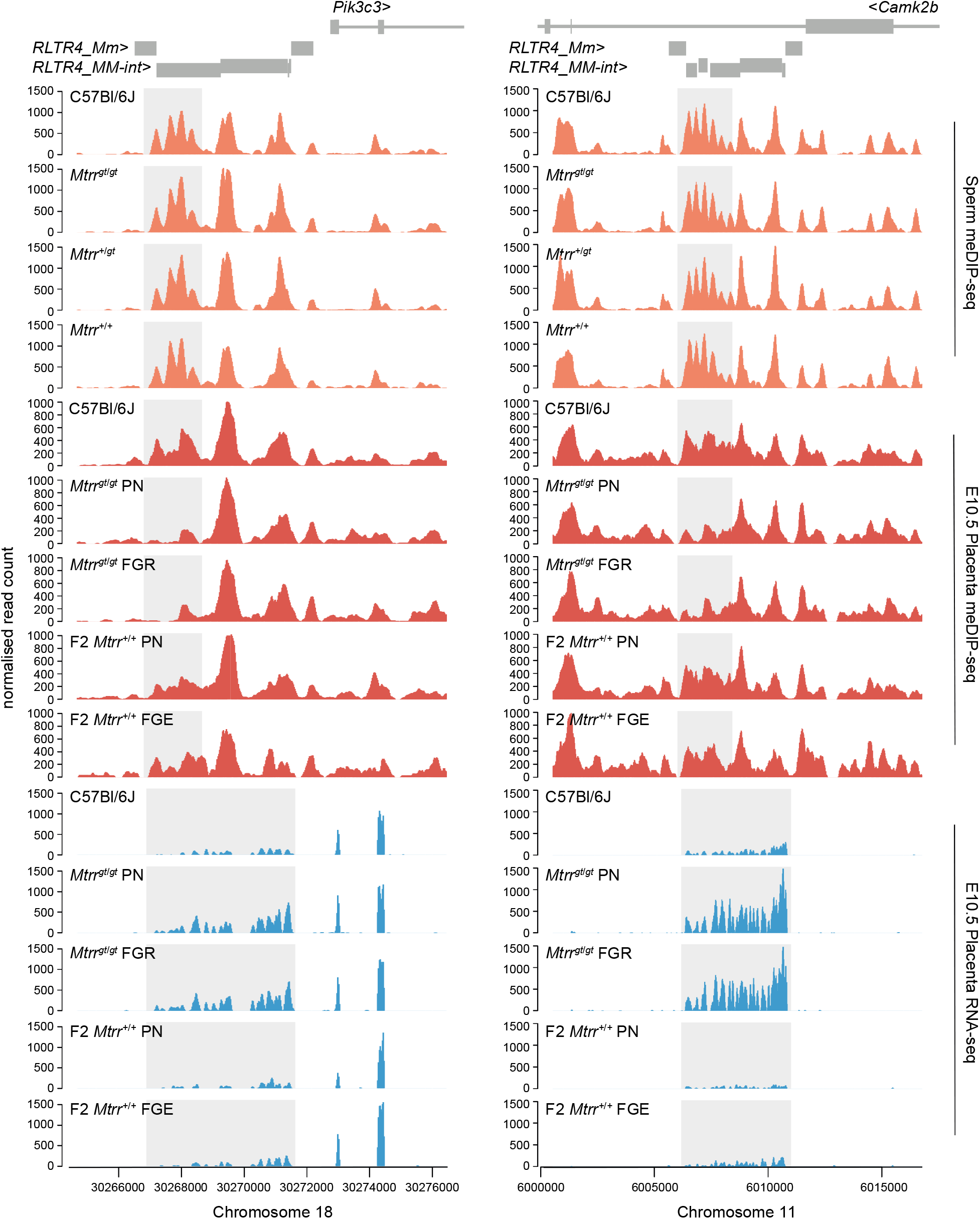
Analysis of sperm and placenta DNA methylation and transcript expression of specific RLTR4 elements. Data tracks showing normalised meDIP-seq and RNA-seq reads across full-length ERVs comprising *RLTR4_Mm* and *RLTR4_MM-int* elements on mouse chromosome 18 associated with the *Pik3c3* gene (left-hand panel) and chromosome 11 associated with the *Camk2b* gene (right-hand panel). Sperm MeDIP-seq data was from adult sperm of C57Bl/6J, *Mtrr^gt/gt^*, *Mtrr^+/gt^*, and *Mtrr^+/+^* mice (orange; N=8 males per experimental group). Placenta meDIP-seq (red) and RNA-seq (blue) data was from placentas at E10.5 of C57Bl/6J conceptuses (N=8 or 6, respectively), *Mtrr^gt^*^/*gt*^ conceptuses associated with phenotypically normal (PN; N=7 or 14, respectively) or fetal growth restricted (FGR; N=7) fetuses, and F2 *Mtrr^+/+^* conceptuses associated with PN (N=8 or 4, respectively) or fetal growth enhanced (FGE; N=3 or 4, respectively) fetuses. Genomic locations of the DMRs and differential transcript expression is highlighted in light grey.

